# Locating a transgene integration site by nanopore sequencing

**DOI:** 10.1101/520627

**Authors:** Peter K. Nicholls, Daniel W. Bellott, Ting-Jan Cho, Tatyana Pyntikova, David C. Page

## Abstract

The introduction of foreign DNA into cells and organisms has facilitated much of modern biological research, and it promises to become equally important in clinical practice. Locating sites of foreign DNA incorporation in mammalian genomes has proven burdensome, so the genomic location of most transgenes remains unknown. To address this challenge, we applied nanopore sequencing in search of the site of integration of *Tg(Pou5f1-EGFP)^2Mnm^* (also known as *Oct4:EGFP*), a widely used fluorescent reporter in mouse germ line research. Using this nanopore-based approach, we identified the site of *Oct4:EGFP* transgene integration near the telomere of Chromosome 9. This methodology simultaneously yielded an estimate of transgene copy number, provided direct evidence of transgene inversions, revealed contaminating *E. coli* genomic DNA within the transgene array, validated the integrity of neighboring genes, and enabled definitive genotyping. We suggest that such an approach provides a rapid, cost-effective method for identifying and analyzing transgene integration sites.

## Introduction

The experimental integration of foreign DNA sequences into mammalian genomes has generated a wide range of genetic tools for basic research and facilitated fundamental discoveries that have revolutionized biology and medicine (Jaenisch and Mintz 1974). In mice, injection of foreign DNA into zygotes is commonly used to produce transgenic offspring that carry gain-of-function alleles, reporter alleles, or express specialized enzymes, such as Cre recombinase for conditional gene deletion. Such transgenes typically integrate at random positions within the genome, and not in single copy but as tandemly-arranged copies, and they may disrupt the function of endogenous genes at or near the site of transgene integration (Dennis *et al*. 2012; Laboulaye *et al*. 2018; Goodwin 2017). It is therefore critical to determine the site of transgene integration to ensure that the function of endogenous genes has not been disrupted. Identifying sites of transgene integration also permits the development of PCR assays that distinguish between offspring that are heterozygous or homozygous for the transgene locus.

Current methods to identify sites of integration are labor intensive or require the design of specialized assays and computational expertise (Itoh *et al*. 2015; Cain-Hom *et al*. 2017). For these reasons, only 433 of the 8,715 transgenic alleles reported in the mouse genome database are currently annotated with a chromosomal location (www.informatics.jax.org/).

To overcome these challenges, we applied nanopore sequencing in search of the genomic site of integration of the *Tg(Pou5f1-EGFP)^2Mnm^* transgene (*Oct4:EGFP*), a widely used fluorescent reporter allele that expresses an EGFP under the control of the distal enhancer of *Pou5f1* (Szabó *et al*. 2002).

## Results

We isolated high-molecular-weight DNA fragments from an adult male mouse carrying the *Oct4:EGFP* allele, and applied this to a MinION flow cell (Oxford Nanopore Technologies, Oxford UK), a sequencing platform capable of producing ultra-long sequence reads of individual DNA strands. From eight sequencing runs, we obtained 1.8-fold haploid genome coverage. We aligned these reads to the nucleotide sequence that encodes EGFP, as well as to the reference genome of the mouse. We identified 25 reads that contained one or more copies of EGFP; all of these also contained sequences that matched the promoter and coding region of the *Pou5f1* gene on chromosome 17, as expected for reads derived from the transgene locus. By constructing a minimal tiling path, we found direct nanopore sequencing evidence of at least six copies of the *Oct4:EGFP* transgene. From a comparison of coverage depth, we estimated that the transgene is present in about 18 copies per allele. Most copies are arranged in tandem, but we found nanopore evidence for at least one change in the direction of the array, captured in two separate nanopore reads (Figure 1A,B). To quantify the number of *EGFP* copies per transgene allele, we performed qPCR on genomic DNA isolated from mice heterozygous, or homozygous, for the *Oct4:EGFP* transgene. As a reference, we normalized *GFP* detection to genomic DNA from mice that carry a single copy of *GFP* inserted at the *Nanog* locus (*1xGFP*). We found that each *Oct4:EGFP* allele carries 25.9 + 2.4 copies of *EGFP* (mean, SD, Figure 1C), representing a total transgene insertion of approximately 450 kb. We also obtained two reads that terminated with a sequence that aligned to the *E. coli* DH5α genome, suggesting contamination with bacterial DNA during production of the transgenic mouse line (Figure 1D). By PCR, we determined that the total length of this *E. coli* sequence is 6.2 kb, and is followed by a transgene copy in the opposite orientation, representing a second inversion in the transgene array.

**Figure 1.**
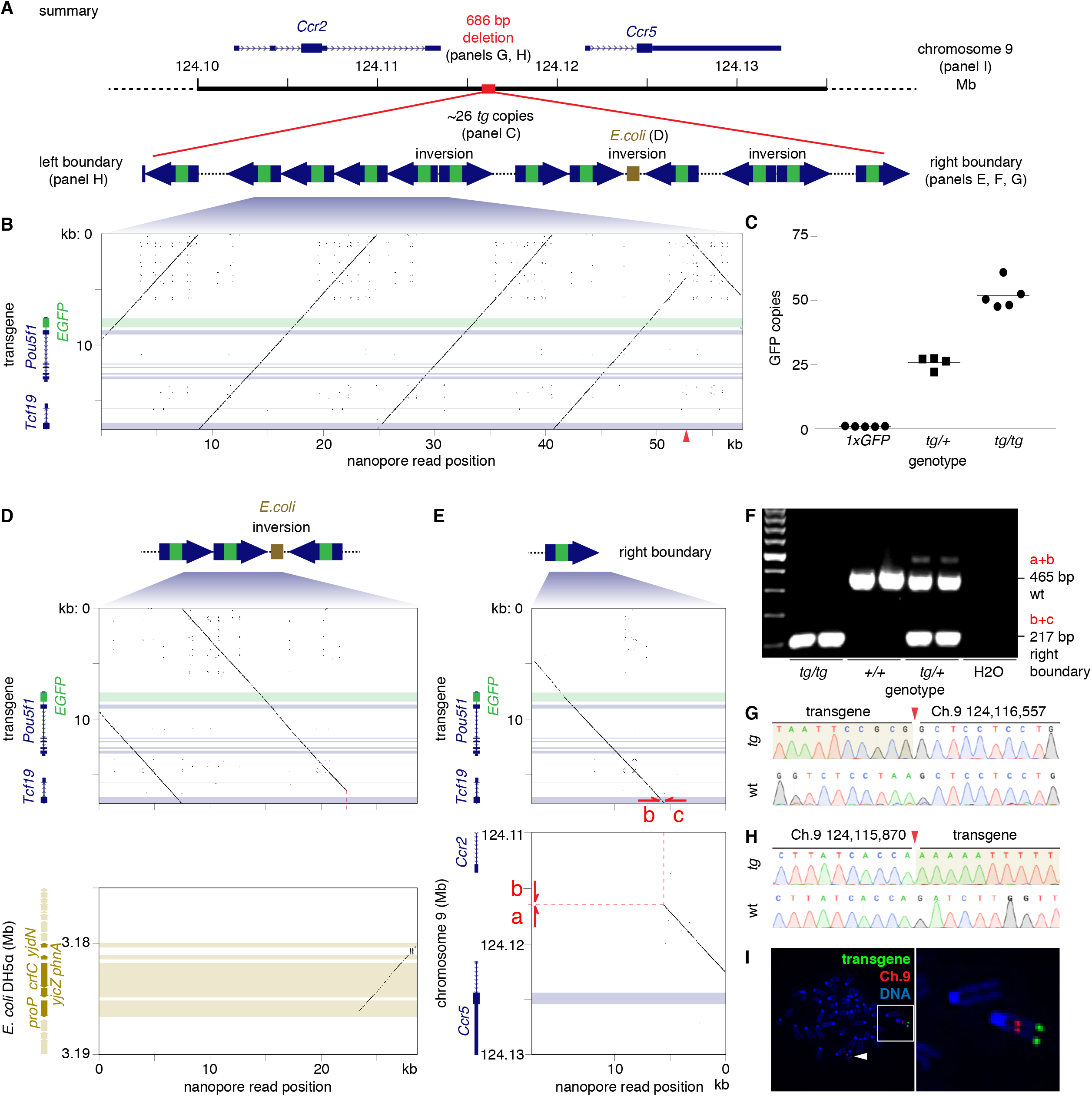
**(A)** Overview of *Oct4:EGFP* transgene structure, with call-outs to subsequent panels in this figure. **(B)** Nucleotide dot plot; all plots with window size 15 nucleotides and step size one nucleotide; horizontal bars indicate coding sequences. 57.7 kb nanopore read (x-axis) compared with predicted insert sequence of transgene, containing distal enhancer and *Pou5f1* CDS with *EGFP* reporter in first exon (y-axis). Red arrow-head on x-axis marks an inversion. **(C)** Quantitative PCR of *GFP* indicates that the transgene is present in about 26 copies per *Oct4:EGFP* allele. Nucleotide dot plots of **(D)** nanopore read aligning to two transgene copies (upper panel) and *E. coli* DH5α genome (lower panel) and **(E)** nanopore read containing 13 kb of transgene insert (upper), and 5.5 kb region between *Ccr2* and *Ccr5* on chromosome 9 (lower). Dashed red line marks *Oct4:EGFP* integration site. Red arrows mark PCR primers flanking the insertion site that **(F)** produce a PCR product of expected size according to the mouse genotype. **(G and H)** Sequencing transgene boundaries confirms transgene integration into sub-telomeric region of chromosome 9 (red arrow-head). *tg: Oct4:EGFP* transgene, wt: wild-type. **(I)** FISH on mouse embryonic fibroblast cell line, heterozygous for the transgene, hybridized with probes for *Oct4:EGFP* transgene (green) and Chromosome 9 (red). DNA stained with DAPI (blue). Inset: transgene at distal telomere of Chromosome 9, white arrow-head: wild-type Chromosome 9.

Of the 25 nanopore reads derived from the transgene locus, one read also contained a 5.5 kb sequence mapping to mouse Chromosome 9 in an intergenic region between *Ccr2* and *Ccr5* (Figure 1E). These observations suggested that the *Oct4:EGFP* transgene is located in the sub-telomeric region of Chromosome 9.

To confirm transgene integration between *Ccr2* and Ccr5, we designed primers spanning the right boundary, between this region on Chromosome 9 and the *Oct4:EGFP* transgene sequence, and identified a single PCR product that was observed with mice carrying the *Oct4:EGFP* transgene, but not in control mice (Figure 1F). Sanger sequencing confirmed integration of the *Oct4:EGFP* transgene at Chromosome 9:124,116,557 in the reference genome (Figure 1G). Using primers designed to amplify the reference DNA sequence spanning the putative integration site, a PCR product was detected only in mice carrying at least one copy of the reference wild type allele (Figure 1F).

To identify the left boundary of the transgene integration, we designed PCR primers adjacent to the 3’UTR of *Ccr2* and performed long-range PCR with a primer to exon 3 of *Tcf19* in the *Oct4:EGFP* transgene sequence. By Sanger sequencing of the resulting PCR product, we identified the left boundary of integration at Chromosome 9:124,115,870 (Figure 1H), followed by a sequence aligning to the distal enhancer of *Pou5f1* starting at 17:35,501,352, and running in the reverse orientation. Collectively, these findings indicate that integration of the *Oct4:EGFP* transgene into Chromosome 9 was accompanied by a deletion of 686 bp of the reference sequence between *Ccr2* and Ccr5, and the incorporation of approximately 26 copies of the *EGFP*, representing the insertion of approximately 450 kb of DNA (Figure 1A). To validate our findings and confirm transgene integration on Chromosome 9, we derived somatic feeder cells from embryos heterozygous for the *Oct4:EGFP* transgene, and conducted fluorescence *in situ* hybridization (FISH). As expected, we observed a single hybridization site near the telomere of Chromosome 9 (Figure 1I).

Finally, we determined whether protein coding genes surrounding the site of *Oct4:EGFP* transgene integration remained intact. For each gene encoded by the reference contig GL456153.2 (*Ccr1, Ccr1l, Ccr3, Ccr2* and Ccr5), we obtained at least one nanopore read encompassing the entire protein coding sequence, confirming that each gene is present in the genome of *Oct4:EGFP* mice (Figure 2). To fill the remaining gaps present in the gene bodies of *Ccr3* and *Ccr2*, we designed PCR assays extending beyond the nanopore sequencing reads, and detected products of the correct size in mice homozygous for the *Oct4:EGFP* transgene (Figure 2, Supplemental Table 2). Sanger sequencing confirmed that these PCR products matched the expected sequence (not shown), indicating that genes within this contig were unaffected by integration of the transgene.

**Figure 2.**
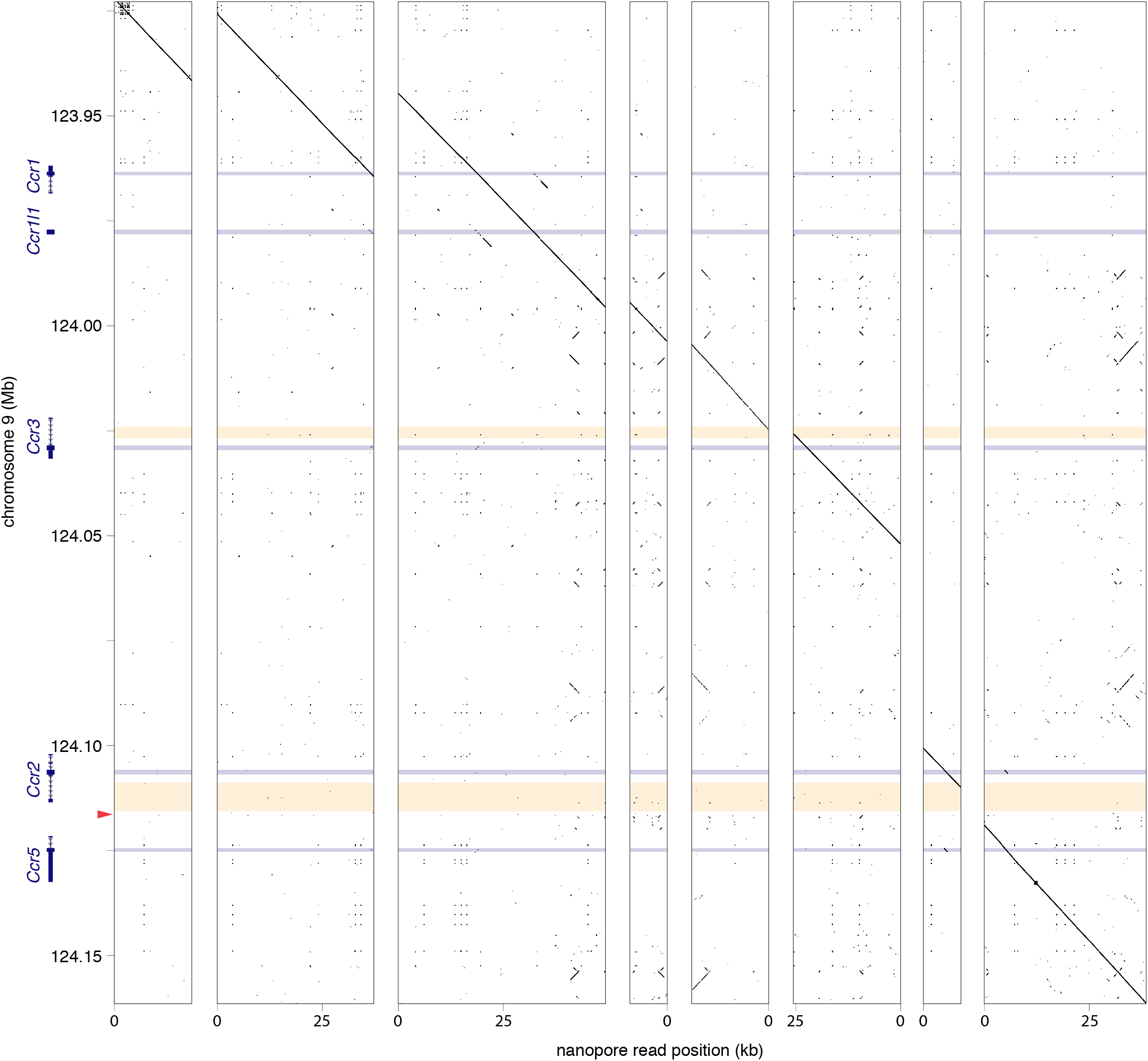
Nucleotide dot plots with window size 15 nucleotides and step size one nucleotide. Blue horizontal bars: protein coding sequences; yellow horizontal bars: intragenic regions spanned by PCR assays (Supplemental Table 2); red arrowhead: *Oct4:EGFP* transgene integration site. Eight nanopore reads mapping to contig GL456153.2 on Chromosome 9. All genes adjacent to the transgene integration site (*Ccr1, Ccr1l, Ccr3, Ccr2*, and Ccr5) were present in our reads, and not disrupted by transgene insertion.

## Discussion

The advent of whole-genome ultra-long-read sequencing technologies provides a platform for the rapid and cost-efficient discovery of transgene integration sites. By applying this technology to the mouse genome, we identified the location of the widely used *Oct4:EGFP* reporter allele, and established a genotyping protocol to distinguish between mice heterozygous or homozygous for the transgene. In addition to identifying the boundaries of transgene integration, ultra-long sequencing reads simultaneously enable an assessment of genome integrity, an estimate of copy number, and the detection of contaminating DNA at the site of transgene integration. We suggest this approach can readily be applied to other transgenic organisms or cell lines (Tyson *et al*. 2018), where identification or validation of transgene integration sites may prove critical, such as prior to the application of cell therapy (Dunbar *et al*. 2018).

## Acknowledgements

*B6-Nanog^tm1Hoch^* and 129S4/SvJae mice were a gift from Rudolf Jaenisch. This work was supported by the Howard Hughes Medical Institute, where DCP is an Investigator. PKN is a recipient of the Hope Funds for Cancer Research Fellowship (HFCR-15-06-06). The funders had no role in study design, data collection and analysis, decision to publish, or preparation of the manuscript.

## Author Contributions

PKN and DWB conceptualized the project and designed the experiments. PKN maintained the mouse lines, collected mouse tissue samples, and performed PCR. DWB analyzed nanopore sequencing reads and performed all bioinformatics analyses. TJC isolated high molecular weight DNA and operated the nanopore. TP cultured MEF cells and performed FISH. PKN, DWB and DCP wrote the paper. DCP supervised the project.

## Author Information

The authors have declared that no competing interests exist. Correspondence and requests for materials and methods should be addressed to.

## Materials and Methods

### Mouse maintenance

All experiments involving mice conformed to ethical principles and guidelines approved by the Committee on Animal Care at the Massachusetts Institute of Technology. Mice homozygous for the germ line reporter allele *CBA-Tg(Pou5f1-EGFP)^2Mnn^ (Oct4:EGFP*) (Szabó *et al*. 2002), and also carrying the *Sry^tm1^* deletion (Wang *et al*. 2013) and *Tg(Sry)2Ei* transgene (Washburn *et al*. 2001) were maintained on a B6 background (Jackson Laboratory, Bar Harbor ME, stock numbers 004654, 010905, and 002994, respectively). The *B6-Nanog^tm1Hoch^ (1xGFP*) reporter allele (Maherali *et al*. 2007) was backcrossed to B6 for 10 generations (Jackson Laboratory, stock number 016233). Wild-type 129S4/SvJae mice were a gift from Rudolf Jaenisch.

### Genotyping and PCR

A small ear biopsy was taken prior to weaning. Genomic DNA was extracted in tissue lysis buffer (100 mM Tris pH 8.5, 5 mM EDTA, 0.2% SDS, 200 mM NaCl, and 100 μg/ml Proteinase K) at 65° overnight. DNA was precipitated with an equal volume of isopropanol and centrifuged. The pellet was then washed in 70% v/v ethanol, centrifuged, and re-suspended in TE buffer (10 mM Tris pH 8.0, 1 mM EDTA). Genotyping was performed using Phusion DNA polymerase (New England Biolabs Inc, Ipswich MA) with the primers listed in Supplemental Table 1. Sanger sequencing of PCR products was performed under contract by Eton Bioscience (Boston, MA), and trace data visualized using SnapGene Viewer software (v3.0.2, GSL Biotech, LLC., Chicago IL). Long range PCR was performed using Advantage 2 Polymerase following the manufacturer’s protocol (Clontech Laboratories, Mountatin View CA).

### Quantitative PCR

We diluted mouse genomic DNA to 5 ng/ul and performed quantitative PCR using an Applied Biosystems 7500 Fast Real-Time PCR instrument and Power SYBR Green PCR Master Mix (Applied Biosystems) with the following cycling conditions: 50° for 2 min, 95° for 10 min, then 95° for 15 s, 60° for 1 min, 75° for 30 s with fluorescent read at 75°, repeated 40 times. To account for total DNA input, we first normalized the relative quantitation of *EGFP* to *Gapdh*. Next, we normalized this ratio to the ratio in mice known to harbor a single copy of *GFP* in the *Nanog* locus (*1xGFP*), and calculated the relative number of *EGFP* copies per transgene.

### Extraction of high molecular weight DNA

We flash froze adult mouse liver in liquid nitrogen, and ground it to a fine powder by mortar and pestle. We then extracted DNA using a modified protocol from Sambrook and Russell, optimized for ultra-long fragments (Jain *et al*. 2018). Briefly, we re-suspended the powder in 10 ml tissue lysis buffer (100 mM Tris pH 8.5, 5 mM EDTA, 0.2% SDS, 200 mM NaCl, and 100 μg/ml Proteinase K) and 100 μL RNase A (20 mg/ml, ThermoFisher Scientific). We extracted protein from the lysate with 10 ml BioUltra TE-saturated phenol (#77607, MilliporeSigma, Burlington MA) using phase-lock gel falcon tubes, followed by extraction of DNA in chloroform-isoamyl alcohol 24:1 (#25666, MilliporeSigma). We precipitated DNA in 4 ml 5 M ammonium acetate mixed with 30 mL ice-cold absolute ethanol, recovered it with a glass hook, and washed it twice in 70% v/v ethanol. Subsequently, we air-dried the DNA pellet and then re-suspended it in 300 μl of 10 mM Tris-HCl pH 8.5, incubated without mixing for 2 days at 4°.

### Nanopore library preparation

For ultra-long reads, we used a modified version of the Rapid Adapters protocol for genomic DNA (SQK-RAD004 Rapid Sequencing Kit, ONT) (Jain *et al*. 2018). Briefly, we combined 16 μl of genomic DNA with 5 μl FRM and mixed slowly by pipetting with a cut-off pipette tip. We then incubated the mixture in a thermocycler at 30° for 75 s, then 75° for 75 s. After this, we added 1 μl RAD and 1 μl Blunt/TA ligase and incubated the ligation at room temperature for 30 min to produce a sequencing library. We diluted the library with a fresh mixture of 25.5 ul Running Buffer with Fuel mix and 27.5 μl NFW. To load the library onto the MinION, we aspirated the 75 μl library with a wide bore tip and slowly pipetted the diluted library onto the SpotOn port as it gets siphoned into the flow cell. We sealed the SpotOn port and equilibrated the flow cell for 45 min prior to the start of the sequencing run to allow long fragments to diffuse into the pores.

### Alignment and data processing

We used Albacore (version 2.3.1, Oxford Nanopore Technologies) to transform raw nanopore fast5 data to basecalls and quality scores. We obtained 611,279 reads with an n50 of 28,297 bp, totaling 4.88 Gbp of sequence across eight runs, or approximately 1.8-fold coverage of the mouse genome. We used minimap2 (Li 2018) (version 2.14-r883) with the ‘-x map-ont’ option to align the fastq sequences of the nanopore reads to the fasta sequences of the mouse reference genome (mm10/GRCm38), the genome of *E. coli* DH5α (GenBank: CP025520.1), and the nucleotide sequence that encodes EGFP (GenBank: U55761.1). We identified 25 reads with one or more matches to EGFP. We counted a total of 32 matches to EGFP in these reads; dividing by 1.8-fold coverage of the genome, we estimated that there were approximately 18 copies of the transgene. After locating the transgene boundary on chromosome 9 in contig GL456153.2, we examined the reads that align to this contig to verify that *Ccr1, Ccr1l, Ccr3, Ccr2*, and *Ccr5* were present in our reads, and not disrupted by the transgene insertion. We generated dot plots using a custom perl script (http://pagelab.wi.mit.edu/material-request.html) and the sequence of GOF18ΔPE EGFP (Addgene plasmid #52382; http://n2t.net/addgene:52382; RRID:Addgene_52382), edited to remove the neomycin resistance cassette and pBluescript KS vector sequence. We obtained gene tracks of the Basic Gene Annotation Set (Ensembl 93) using the UCSC genome browser (mm10/GRC38). We designed primers using primer3 (Untergasser *et al*. 2012).

### Mouse embryonic fibroblast (MEF) derivation and FISH analyses

Primary MEFs were prepared from F1 E13.5 mouse embryos (129S4 x C57BL/6N, where the father was homozygous for the *Oct4:EGFP* transgene). Briefly, the embryo body cavity was minced using sterile blades, and a single cell suspension obtained using 0.25% Trypsin for 15 min. The cell mix was then thoroughly dissociated in MEF media, consisting of DMEM with 10% fetal calf serum, 0.1 M nonessential amino acids, 2 mM L-glutamine, and penicillin/streptomycin (each from Thermo Fisher Scientific, Waltham MA). Cells were then cultured until confluent.

FISH probes were generated from a BAC specific to Chromosome 9 (RP24-402I19), and to a plasmid containing the *Oct4:EGFP* transgene (GOF18ΔPE EGFP, Addgene plasmid #52382; http://n2t.net/addgene:52382; RRID:Addgene_52382).

FISH staining was performed on MEF cells as previously described (Saxena *et al*. 2000). Cells were harvested by trypsinization, treated with 0.075 M KCl hypotonic solution and fixed in methanol:acetic acid (3:1 v/v). The cell suspension was applied to slides and treated with RNase. After dehydration in ethanol, slides were incubated at 80° for 5 min to denature the DNA and washed again in ethanol.

BAC probes were labeled by nick translation with direct label Cy-3 and Biotin (subsequently detected with a fluorescent antibody). The probes were denatured together with unlabeled competitor DNA and applied to the slides. Slides with probes were incubated at 37° for 16 h, washed in 50% formamide to remove non-specific hybridization and reduce background, and dehydrated in ethanol. Chromosomes were stained with DAPI and imaged on a fluorescence microscope.

### Data availability

Data generated by nanopore sequencing and Sanger sequencing has been deposited with GenBank under accession numbers PRJNA508423, MK327979, MK327980, and MK344432. The authors affirm that all other data necessary for confirming the conclusions of the article are present within the article, figures and tables.

